# Recombinant annexin A6 promotes membrane repair in a stem cell derived-cardiomyocyte model of dystrophic cardiomyopathy

**DOI:** 10.1101/2022.03.09.483528

**Authors:** Dominic E. Fullenkamp, Alexander B. Willis, Jodi L. Curtin, Ansel P. Amaral, Sloane I. Harris, Paul W. Burridge, Alexis R. Demonbreun, Elizabeth M. McNally

## Abstract

Heart failure is a major source of mortality in Duchenne muscular dystrophy (DMD). DMD arises from mutations that ablate expression of the protein dystrophin, which render the plasma membrane unusually fragile and prone to disruption. In DMD patients, repeated mechanical stress leads to membrane damage and cardiomyocyte loss. Induced pluripotent stem cell-derived cardiomyocytes (iPSC-CMs) offer the opportunity to study specific mutations in the context of a human cell, but these models can be improved by adding physiologic stressors. We modeled the primary defect underlying DMD by applying equibiaxial mechanical strain to DMD iPSC-CMs. DMD iPSC-CMs demonstrated an increased susceptibility to equibiaxial strain after 2 hours at 10% strain relative to healthy control cells, measured as increased lactate dehydrogenase (LDH) release. After 24 hours, both DMD and healthy control iPSC-CMs showed evidence of injury with release of LDH and cardiac troponin T. We exposed iPSC-CMs to recombinant annexin A6, a protein resealing agent, and found reduced LDH and troponin release in DMD and control iPSC-CMs that had been subjected to 24 hour strain at 10%. We used aptamer protein profiling of media collected from DMD and control iPSC-CMs and compared these results to serum protein profiling from DMD patients. We found a strong correlation between the proteins in DMD patient serum and media from DMD iPSC-CMs subjected to mechanical stress. By developing an injury assay that specifically targets an underlying mechanism of injury seen in DMD-related cardiomyopathy, we demonstrated the potential therapeutic efficacy of the protein membrane resealer, recombinant annexin A6, for the treatment of DMD-related cardiomyopathy and general cardiac injury.

## Introduction

Duchenne muscular dystrophy (DMD) is an X-linked disease that results from mutations in the *DMD* gene, which codes for the protein dystrophin (1). Clinically, DMD presents in the first decade with weakness and markedly elevated serum biomarkers like creatine kinase (CK) (2). Cardiac involvement is typically evident by the second decade and contributes to morbidity and mortality in DMD (3). In heart and muscle, dystrophin localizes to the plasma membrane and is concentrated in the membrane above the Z-disc, colocalizing with other proteins of the dystrophin complex, including the sarcoglycans and dystroglycans (4–6). This complex forms a critical transmembrane structural and signaling connection between the sarcomere and the extracellular matrix (4,7–10). Disruptions along this axis produce membrane fragility and account for multiple forms of muscular dystrophy with cardiac involvement (11–13). Daily glucocorticoid administration, begun in the mid-first decade, delays loss of mobility by ~2 years (14). Early initiation of ACE inhibitors slows progression of cardiomyopathy (15–17), and cardiomyopathy treatment and heart failure management in DMD largely relies on guideline-directed heart failure strategies (3,18). Several antisense-mediated exon skipping agents are now approved for use in DMD, but these agents have relatively poor penetration into the myocardium and are useful for less than 25% of DMD mutations (19,20). Novel therapeutics for the treatment of DMD are currently under investigation, including gene replacement therapy with micro-dystrophins, gene editing approaches, and membrane re-sealants (21–24). For clinical agents treating skeletal muscle in DMD, most studies have relied on endpoints like time to loss of mobility or measures of muscle strength or performance. Clinical trials for DMD cardiomyopathy are complicated by patients having reduced or no ambulatory capabilities (20).

iPSC-derived cardiomyocytes (iPSC-CMs) can be used to evaluate patient-specific therapies in a human cell context (25). However, iPSC-CMs are immature in nature and are generally cultured under conditions that fail to mimic the load and stress seen by the human heart (26,27). Despite progress with tissue engineering methods, which can partially improve maturity (28,29), approaches to evaluate dynamic physiologic mechanical stress are still under development. Studies using rat neonatal cardiomyocytes or mouse embryonic fibroblasts investigated the effects of mechanical stress to understand early signaling responses that lead to cardiac hypertrophy (30) and pathological signaling responses in nuclear membrane defects (31). We now investigated the differential response of mechanical stress on DMD and healthy control iPSC-CMs and the response to a resealing protein, recombinant annexin A6, which was previously identified as a genetic modifier of muscular dystrophy and a potential therapeutic target (32,33). We further used an aptamer-based protein profiling system to characterize protein release at baseline and in response to mechanical stress responses in DMD iPSC-CMs, finding significant correlation with human serum biomarkers from DMD patients.

## METHODS

### iPSC generation, iPSC culture, cardiac differentiation, enrichment, and expansion

Cells were obtained from a DMD patient and reprogrammed using published methods to generate the cell line DMD-G01 (34). The control line iPSC line (GM033488) was previously described (35). iPSC culture and differentiation were performed per previously published methods (35,36).

### Preparation of flexible membranes and application of equibiaxial strain

Expanded iPSC-CMs were harvested by collagenase digestion as above and plated at a density of 1.5 million cells/well in B27 in RPMI 1640 with 10% FBS (Gibco, 26140079) and 1% penicillin/streptomycin (Gibco, 15070063). Media was exchanged with B27 in RPMI 1640 and 1% penicillin/streptomycin, every-other day. Cyclic sinusoidal equibiaxial strain at 1 Hz was applied using a FX-6000T™ Tension System (FlexCell International).

### Recombinant annexin A6

iPSC-CMs were treated with recombinant annexin A6 (33) at a concentration of 10 μg/mL.

### Biomarker measurement

LDH and cardiac troponin T release was quantified per manufacturer instructions using Promega LDH-Glo™ Cytotoxicity Assay (Promega J2380) and human cardiac troponin T ELISA kit (Abcam, ab223860).

### SOMAscan assay and analysis

The SOMAscan aptamer assay reports 7322 aptamer-based proteomics results per sample in units of Relative Fluorescent Units, which were read into R studio using the *SomalogicsIO* R package (37). The comparison DMD dataset was previously reported (38). Shapiro-Wilk tests were applied to the datasets with an alpha value of 0.05 set as a threshold for normality. Around 40% of both raw and log transformed datasets failed to meet this threshold, therefore non parametric Wilcoxon-Mann-Whitney tests were used to assess differential serum biomarker levels across experimental groups. To account for multiple hypothesis testing, the Benjamini-Hochberg correction method (39) was utilized using the p.adjust function of the *stats* R package (40). Thresholds for significant differential biomarker levels were set at FDR < 0.05 and absolute value of log_2_(fold-change) > 0.5. All statistics were performed in R studio running R version 4.0.2 (2020-06-22) with additional packages (41–44). Volcano plots were generated using R package ggplot2, with the cutoffs for differential levels described above. Heat maps were generated using the R package gplots (45). Gene ontology analysis was performed on differential biomarkers for pathway enrichment using the enrichr online analysis software and the Human WikiPathways 2021 database (46,47).

### Statistical methods

Data was analyzed using Prism 9.3.0. Where comparisons of two conditions were made an unpaired t test was used. Where comparisons of more than two conditions were made an ordinary one-way ANOVA was used with Tukey correction for multiple comparisons. In all cases, p< 0.05 was defined as statistically significant. Statistical data is reported as mean ± SEM. Confidence intervals are reported as 95% (95% CI).

## RESULTS

### Generation, differentiation and expansion of high-quality iPSC-CMs

IPSCs were generated from a DMD patient with an out-of-frame, large deletion spanning *DMD* exons 46-47 (**Figure 1A**). This patient also had a Wolf-Parkinson-White EKG pattern that eventually required ablation with following symptomatic orthodromic reentry tachycardiac localized to the left lateral pathway (**Figure 1B**). The patient had a typical DMD course with loss of ambulation before the age of 11 and developed an associated severe cardiomyopathy with LVEF ~12% despite guideline-directed therapy and biventricular chronic resynchronization therapy (**Figure 1C**). To improve variability in iPSC-CM differentiation, we employed a two-step iPSC-CM enrichment (**Figure 2A**). iPSCs were initially differentiated into iPSC-CMs by conventional methods (35,36), followed by a second step in which iPSC-CMs were enriched using a magnetic separation system. Assessment, pre- and post-enrichment by magnetic separation confirmed improved cardiac troponin T positivity (**Figure 2B** and **2C**). This enriched iPSC-CM cell population was then expanded by adapting a recently published method (48). Combining iPSC-CM enrichment with expansion generated sufficient numbers of high-quality iPSC-CMs for downstream applications.

**Figure 1.**
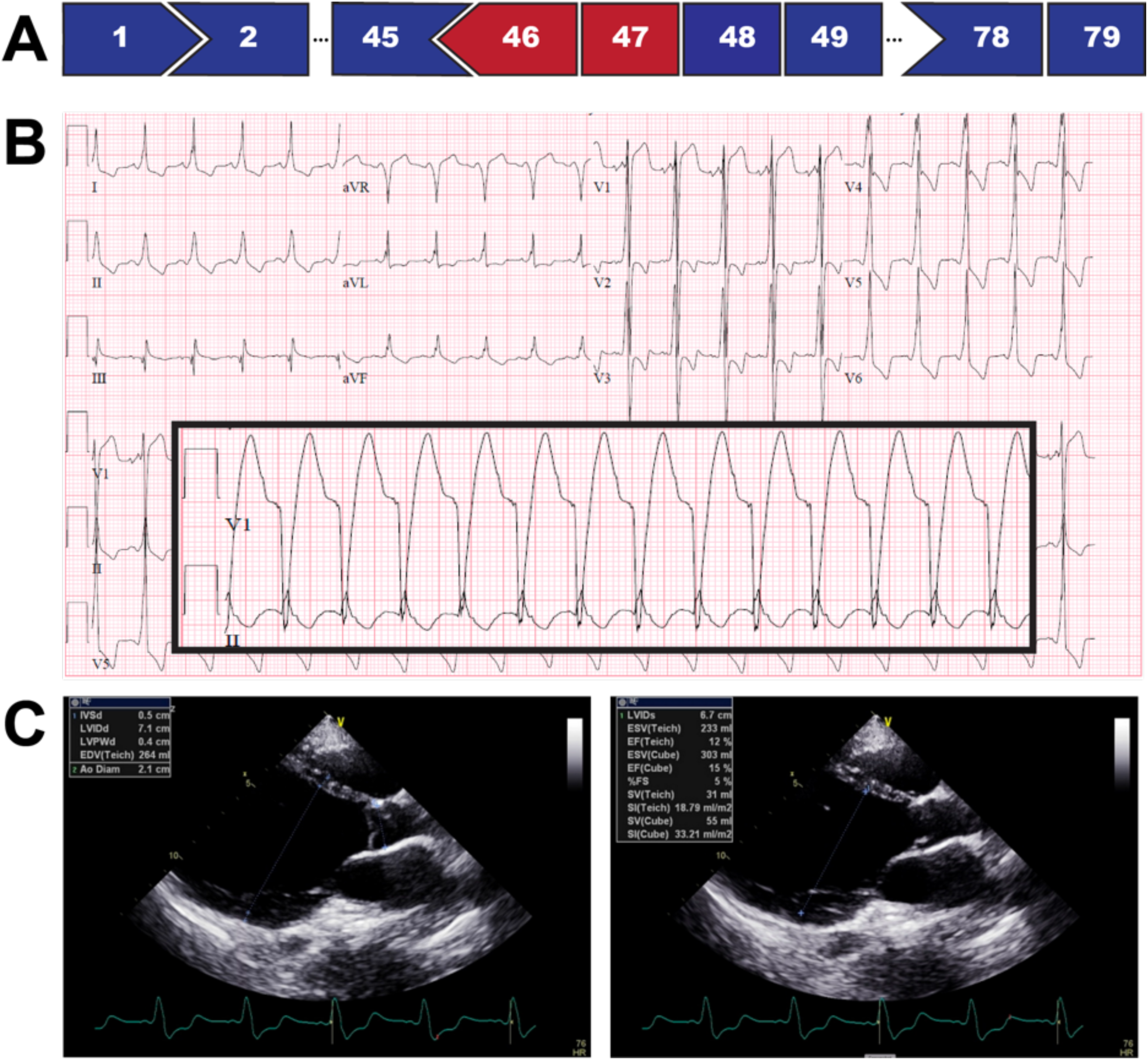
Clinical characteristics of DMD patient. **A**) Abbreviated *DMD* exon map, showing an out of frame exon 46-47 deletion, highlighted in red. **B)** Baseline electrocardiogram, showing a Wolf-Parkinson-White, preexcitation pattern. Inset shows a wide complex tachycardia that was confirmed to be a left lateral pathway orthodromic reentry tachycardia by an electrophysiology study. **C)** Still images from an echocardiogram at age 27, demonstrating an end diastolic dimension of 7.1 cm and end systolic dimension of 6.7 cm. Ejection fraction, by biplane measurement was 12%.

**Figure 2.**
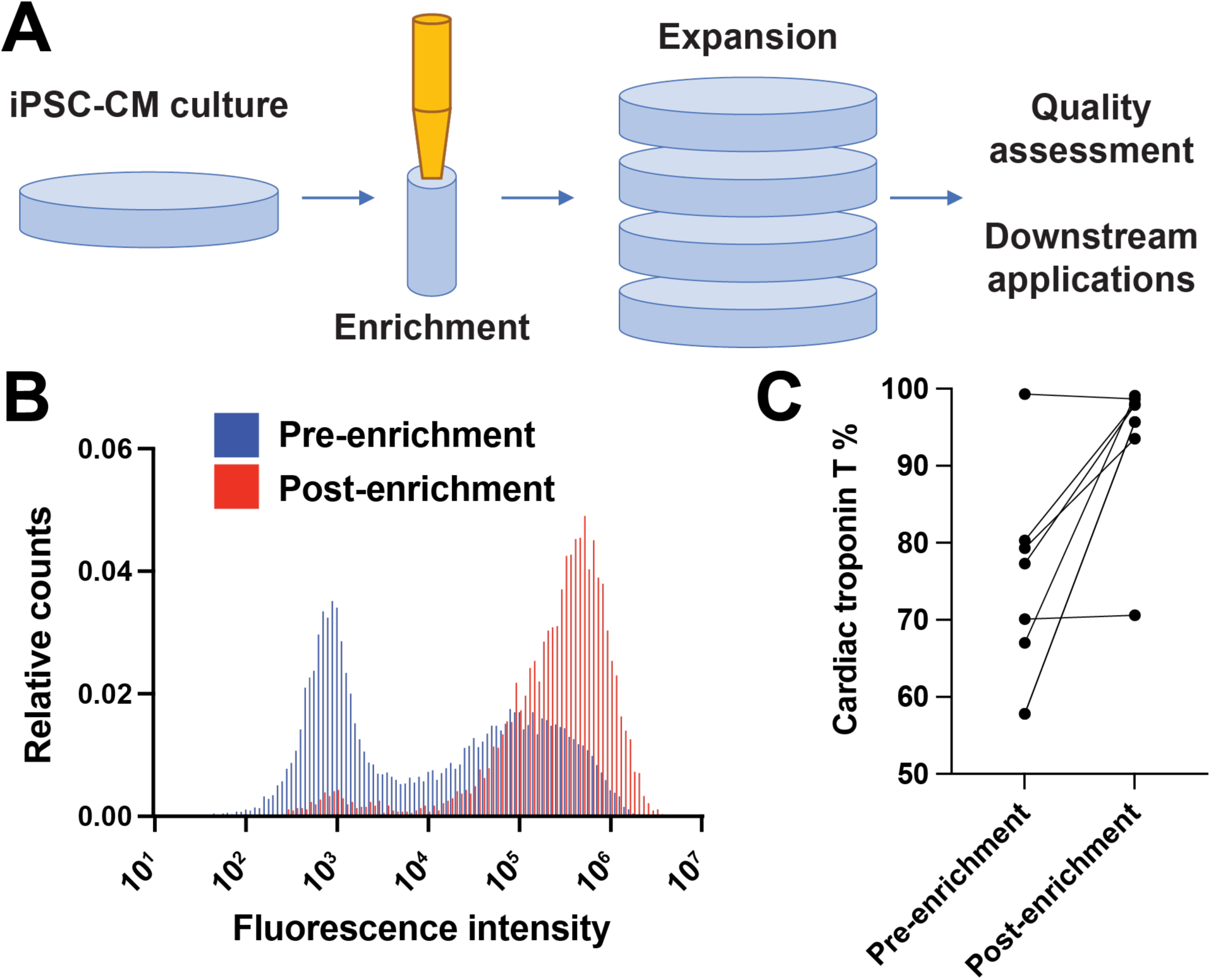
Generation strategy of iPSC-CMs and quality assessment. **A)** Overview of generation, enrichment, and expansion strategy with quality assessment by cardiac troponin T flow cytometry. After differentiation, iPSC-CM are first enriched using the Miltenyi MACs system followed by expansion. **B)** Representative cardiac troponin T staining as assessed by flow cytometry before and after enrichment with increase in cardiac troponin T positivity from 58.7 % to 95.7%. **C)** Validation of enrichment strategy, showing change in cardiac troponin T positivity pre- and post-enrichment for DMD-G01 line.

### Increased susceptibility of dystrophic iPSC-CMs to mechanical strain

Dystrophin deficient cardiomyocytes from animal models have increased susceptibility to mechanical stress relative to controls (21,49). Similarly, serum biomarkers reflective of membrane leak are elevated in DMD patients (38,50). Therefore, we initially sought to define a physiologic degree of mechanical stress to impart on iPSC-CMs that differentiated DMD iPSC-CMs from healthy control iPSC-CMs. iPSC-CMs were plated onto flexible membranes in a 6 well plate format and radial deformation was applied to impart a homogenous equibiaxial strain on plated cells in vitro (**Figure 3A**). Healthy control iPSC-CMs and DMD iPSC-CMs were subjected to 2 h of 0% (unflexed), 5%, 10% or 15% strain and the cell culture media was collected for biomarker determination (**Figure 3B**). Lactate dehydrogenase (LDH) is a clinically relevant serum biomarker of tissue injury, including cardiac injury (51). Control iPSC-CM media LDH levels after 5% and 10% strain remained similar to that of unflexed cells (**Figure 3C**). At 15% strain there was an increase LDH release, however there was also an increase in the variability of the data, likely from the severity of the injury. Media collected from DMD iPSC-CMs showed a dose-dependent increase in LDH levels following strain injury (**Figure 3D**), demonstrating that dystrophic iPSC-CMs are more susceptible to strain-induced injury compared to control iPSC-CMs. Similar to control iPSC-CMs, the variability of LDH release for DMD iPSC-CMs increased at 15% strain, likely related to the severe injury at this high level of strain. Based on our initial considerations to define a physiologic degree of mechanical stress, we observed that 10% strain did not result in significant LDH release in control iPSC-CMs, while it resulted in a significant increase LDH release in DMD iPSC-CMs. Thus, subsequent experiments were performed at 10% strain.

**Figure 3.**
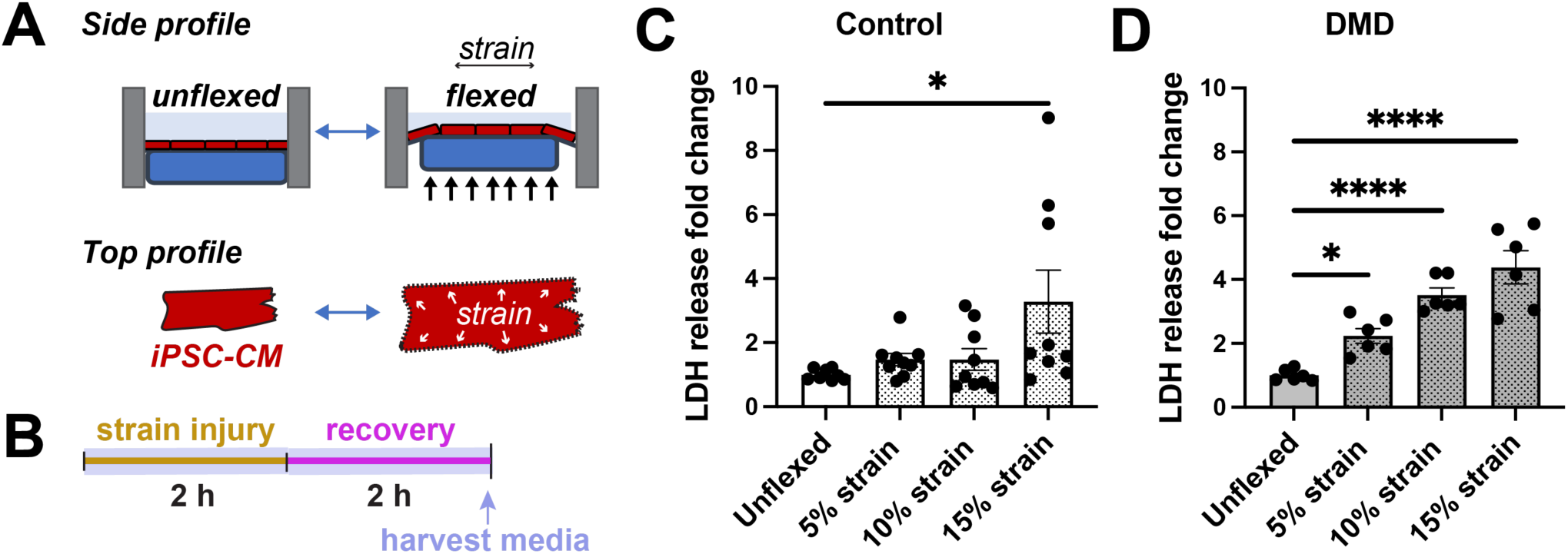
DMD iPSC-CMs show a differential response to equibiaxial strain. **A)** Schematic of application of mechanical stress using the Flexcell system that deforms iPSC-CMs adhered to flexible silicone elastomer membranes using a rigid post, imparting equibiaxial strain. **B)** Overview of injury protocol timeline. iPSC-CMs are subjected to mechanical stress for 2 h followed by a 2 h recovery period. Media is then harvested to determine total LDH release. **(C)** Control iPSC-CMs do not show a significant increase in the release of LDH compared to unflexed conditions at 5% and 10% strain. At 15% strain, LDH fold release increased by 2.3 (95% CI: 0.2 to 4.4, *p = 0.032) (n ≥ 9 from multiple differentiations). **D)** DMD iPSC-CMs show an increase susceptibility to mechanical stress-induced injury compared to healthy control iPSC-CMs. At 5%, 10%, and 15%, LDH fold release increased relative to unflexed conditions by 1.2 (95% CI: 0.02 to 2.5, *p = 0.045), 2.51 (95% CI: 1.3 to 3.7, ****p <0.0001), and 3.4 (95% CI: 2.2 to 4.6, ****p < 0.0001), respectively (n ≥ 6 from multiple differentiations).

### Recombinant annexin A6 promotes membrane repair in control iPSC-CMs

Having defined a strain exposure that differentiated between DMD and healthy control iPSC-CMs, we tested whether applying longer exposure to strain could induce injury in healthy control iPSC-CMs (**Figure 4A**). As shown in **Figure 4B**, LDH release fold change increased by 5.1 (95% CI: 2.9 to 7.2, ****p<0.0001) after 24 h of flexing compared to the non-injury-inducing 2 h time period. This demonstrated that 24 h of 10% strain surpassed the threshold required to induce mechanical membrane injury sufficient for LDH leak in control iPSC-CMs. Recombinant annexin A6 was previously shown to promote resealing in mouse skeletal myofibers injured with a laser (32,33), so we assessed the efficacy of recombinant annexin A6 to enhance repair in cardiomyocytes injury using this mechanical injury model. We first assessed whether fluorescently labelled recombinant annexin A6 bound to control iPSC-CMs after strain exposure. As show in **Figure 4C**, relative mean fluorescent intensity increased by 3.7 (95% CI: 2.5 to 5.0, ****p<0.0001) in treated compared to untreated control iPSC-CMs as assessed by flow cytometry, consistent with recombinant annexin A6-iPSC-CM binding. **Figure 4D** depicts the experimental strategy for assessing membrane repair and response to recombinant annexin A6 in which membrane damage is followed by exposure to recombinant annexin A6 or vehicle, and then strain was continued for 1 h, followed by a 2 h recovery period. In the absence of annexin A6, strain resulted in an increase in LDH release fold change of 1.7 (95% CI: 0.4 to 3.0, **p = 0.01) compared to unflexed controls (**Figure 4E**). When annexin A6 is present during the post-injury recovery period, LDH levels were similar to LDH levels of unflexed cells (95% CI: −1.6 to 1.0, p = 0.79). To corroborate these findings, troponin T release was also measured (**Figure 4F**) and was found to be similarly increased with application of mechanical stress (95% CI: 3.5 to 6.7, ****p<0.0001) and reduce to near baseline levels with recombinant annexin A6 treatment (95% CI: −0.4 to 1.1, p = 0.19). Together these data demonstrate that recombinant annexin A6 can promote repair of injured control iPSC-CMs.

**Figure 4.**
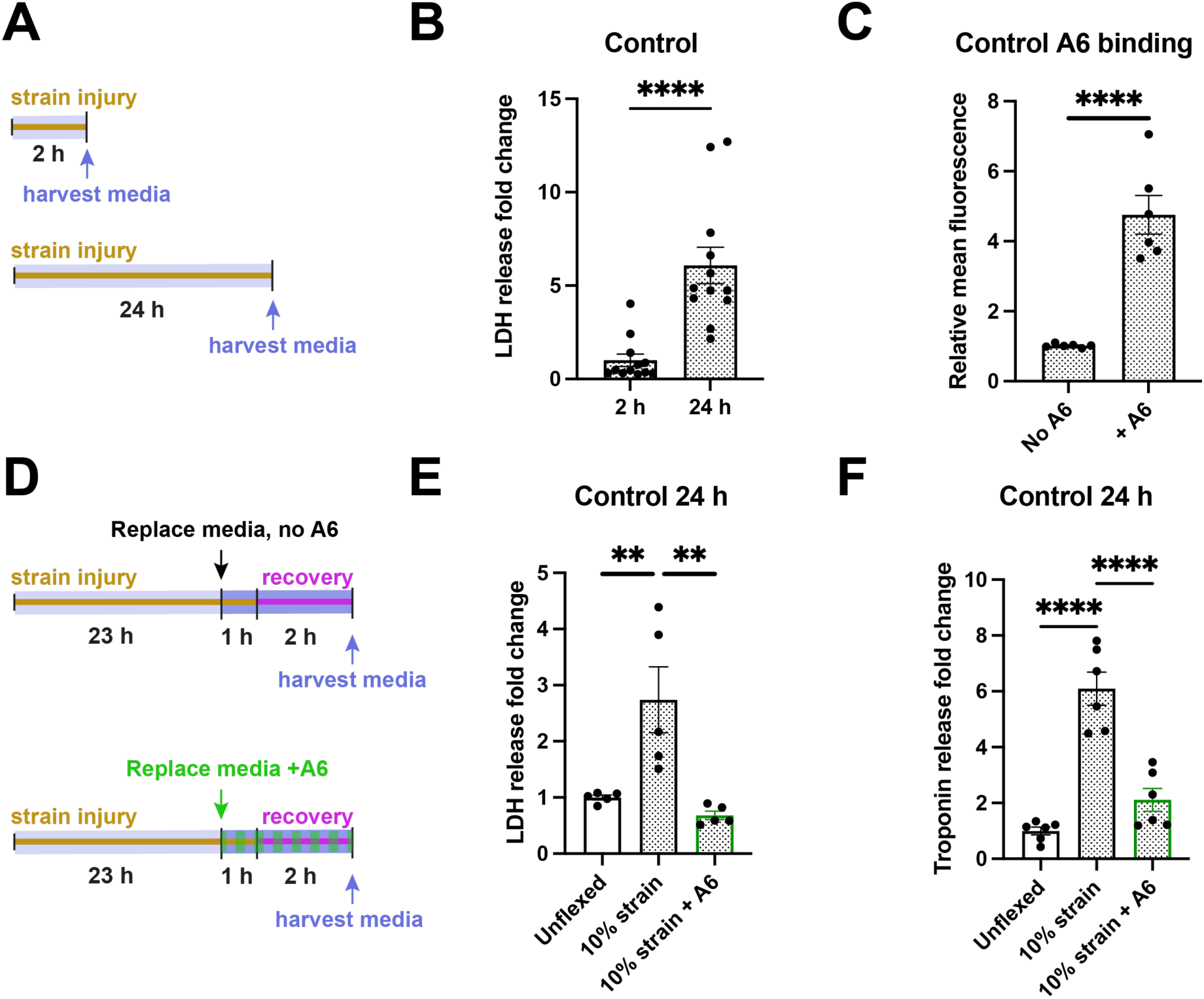
Recombinant annexin A6 enhances repair in healthy control iPSC-CMs. **A)** Schematic of injury protocol comparing 2 h and 24 h at 10% strain since healthy control iPSC-CMs require greater duration of mechanical stress to induce injury. **B)** LDH release fold change increased by 5.1 (95% CI: 2.9 to 7.2, ****p<0.0001) at 24 h compared to 2 h of 10% strain in control iPSC-CMs. n ≥ 12 from multiple differentiations. **C)** Fold change of relative fluorescence intensity increased by 3.7 (95% CI: 2.5 to 5.0, ****p<0.0001) in control iPSC-CMs treated with fluorescently labelled recombinant annexin A6, which was added for the last 1 h of a 10% strain protocol lasting 24 h. n ≥ 6 from multiple differentiations. **D)** Overview of assessment of efficacy of membrane repair for recombinant annexin A6 with a 24 h injury protocol. **E)** LDH release fold change increased by 1.7 (95% CI: 0.4 to 3.0, **p = 0.01) relative to unflexed control iPSC-CMs after a 24 h 10% strain protocol. Recombinant annexin A6 reduced LDH fold release by 2.1 (95% CI: 0.8 to 3.3, **p = 0.003) under a 10% strain protocol relative to untreated strained iPSC-CMs. No significant difference was observed between unflexed and treated 10% strained iPSC-CMs (95% CI: −1.6 to 1.0, p = 0.79). n ≥ 5 from multiple differentiations. **F)** Troponin release fold change increased by 5.1 (95% CI: 3.5 to 6.7, ****p<0.0001) after 10% strain 24 h protocol, while treatment with recombinant annexin A6 under the same protocol reduced troponin release fold change by 4.0 (95% CI: 2.4 to 5.5, ****p<0.0001) with no significant difference compared to the unflexed condition (95% CI: −0.4 to 1.1, p = 0.19) (n ≥ 6 from multiple differentiations).

### Annexin A6 promotes dystrophic iPSC-CM membrane repair

Knowing that dystrophic cells are highly prone to membrane injury, we wanted to determine if recombinant annexin A6 could enhance repair of severely injured DMD iPSC-CMs. We first assessed fluorescently labelled recombinant annexin A6 binding to DMD iPSC-CMs after a 24 h strain protocol. As shown in **Figure 5A**, relative mean fluorescent intensity increased by 3.4 (95% CI: 2.2 to 4.5, ****p<0.0001) in treated strained DMD iPSC-CMs, demonstrating recombinant annexin A6 binding. DMD iPSC-CMs were subjected to the same 24 h, 10% strain injury protocol that is capable of injuring control iPSC-CMs (**Figure 5B**). When recombinant annexin A6 was absent during the post-injury recovery period, LDH release fold change increased by 4.1 (95% CI: 1.2 to 7.0, **p = 0.005) compared to unflexed DMD iPSC-CMs. **Figure 5C** shows that with the addition of recombinant annexin A6 during the post-injury recovery period, LDH levels were similar to unflexed iPSC-CM media (95% CI: −2.8 to 2.9, p = 0.9989) and significantly less than in media from flexed cells lacking annexin A6 (95% CI: 1.2 to 6.9, **p = 0.005). Troponin release fold change mirrored LDH levels, increasing 3.9 (95% CI: 3.1 to 4.7, ****p<0.0001) post-injury in the absence of annexin A6 compared to unflexed controls (**Figure 5D**). Treatment with recombinant annexin A6 reduced fold change troponin levels by 3.5 (95% CI: 2.7 to 4.3, ****p<0.0001) with no significant difference compared to the unflexed condition (95% CI: −0.4 to 1.1, p = 0.52). Consistent with the effects seen on membrane repair in control iPSC-CMs, these results demonstrate efficacy of recombinant annexin A6 in promoting membrane in dystrophic iPSC-CMs.

**Figure 5.**
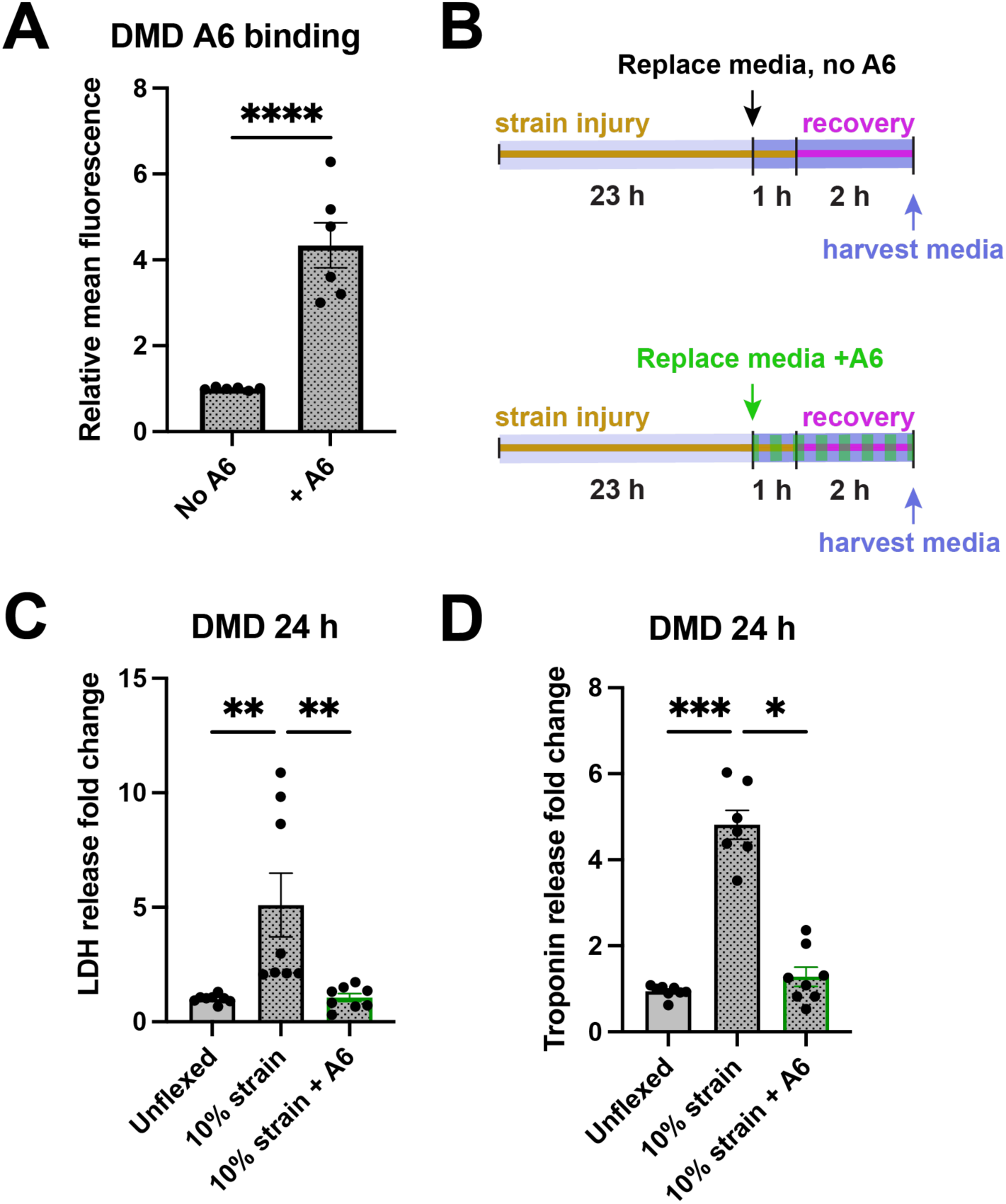
Recombinant annexin A6 enhances repair in DMD iPSC-CMs. **A)** Fold change of mean fluorescence intensity increased by 3.4 (95% CI: 2.2 to 4.5, ****p<0.0001) in DMD iPSC-CMs treated with fluorescently labelled recombinant annexin A6, which was added for the last 1 h of a 10% strain protocol lasting 24 h (n ≥ 6 from multiple differentiations). **B)** Overview of assessment of efficacy of membrane repair for recombinant annexin A6 with a 24 h injury protocol. **C)** LDH release fold change increased by 4.1 (95% CI: 1.2 to 7.0, **p = 0.005) relative to unflexed DMD iPSC-CMs after a 24 h 10% strain protocol. Recombinant annexin A6 reduced LDH release change by 4.0 (95% CI: 1.2 to 6.9, **p = 0.005) under a 10% strain protocol relative to untreated strained iPSC-CMs. No significant difference was observed between unflexed and treated 10% strained iPSC-CMs (95% CI: −2.8 to 2.9, p = 0.9989). n ≥ 8 from multiple differentiations. **D)** Troponin release fold change increased by 3.9 (95% CI: 3.1 to 4.7, ****p<0.0001) after 10% strain 24 h protocol, while treatment with recombinant annexin A6 under the same protocol reduced fold change troponin release by 3.5 (95% CI: 2.7 to 4.3, ****p<0.0001) with no significant difference compared to the unflexed condition (95% CI: −0.4 to 1.1, p = 0.52) (n ≥ 8 from multiple differentiations).

### Stress improves comparison to human biomarkers

A previous study conducted aptamer-based profiling on ambulatory and nonambulatory DMD patients as well as non-dystrophic controls (38). These serum profiles measured 1,125 markers, reflecting both skeletal and cardiac muscle disease in DMD. We employed this same technology to assess biomarker release from flexed and unflexed iPSC-CMs after 2 h at 10% equibiaxial strain. **Figure 6A** shows a volcano plot comparing DMD and control cells in the flexed and unflexed states. Gene ontology analysis of these results demonstrated an increase in matrix metalloproteinase-related proteins at baseline in unflexed DMD cells compared to unflexed control cells (**Figure 6B**). Interestingly flexing caused a response in the aerobic glycolysis pathway in DMD cells relative to control cells (**Figure 6B**). We evaluated clinically the relevant serum biomarkers CKM, LDH, TNNT2, and TNNI3 (**Figure 6C**). Consistent with the results from previous baseline experiments (**Figure 2**), CKM, LDH, and TNNT2, showed similar baseline levels, while TNNI3 had slightly elevated baseline levels in DMD media from iPSC-CMs compared to healthy control. Flexing resulted in a significant increase in CKM, LDH, TNNT2, and TNNI3 in DMD media compared to healthy control, consistent with the clinical phenotype. With respect to matrix metalloproteinases, **Figure 6C** demonstrates an increase TIMP1, TIMP2, MMP2, and MMP9 in unflexed DMD media compared to healthy control media, while only TIMP1 and TIMP2 remained increased relative to control cells as a result of flexing. Therefore, mechanical stress promotes the release of some biomarkers from healthy control cells, consistent with an injury profile.

**Figure 6.**
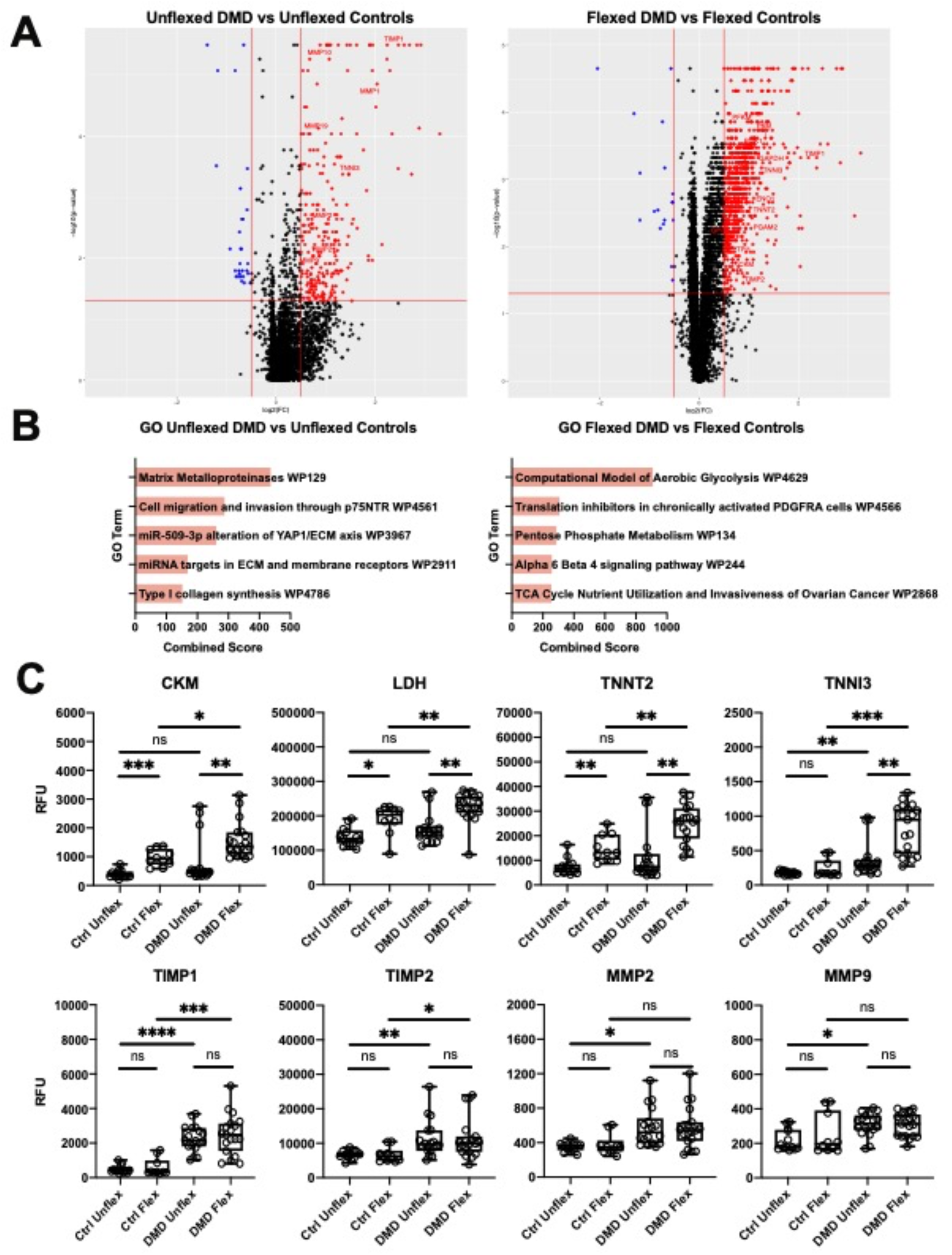
Aptamer protein analysis of media from control and DMD iPSC-CMs. **A)** Volcano plots comparing aptamer profiles from control and DMD iPSC-CMs (left, unflexed; right, flexed). **B)** Corresponding gene ontology analysis of (A). **C)** Specific biomarker data. Top row represents aptamer measurements of are clinically measurements biomarkers. Bottom row represents matrix metalloproteinase relevant proteins.

Hathout, et al. (38) previously defined serum proteins in DMD related to stage of disease. They identified Group 1 proteins as increased in young DMD patients compared to non-dystrophic controls, and these Group 1 markers decreased over time in DMD patients, consistent with loss of muscle mass and reduced locomotion. We expected Group 1 to be most similar to the conditions mimicked by iPSC-CMs, where cells were intact and injured but pathologic fibrosis and myocyte loss were not present. **Figure 7** shows a heatmap of these protein markers. Mechanical stress induces minimal changes in control cells, while mechanical stress strikingly differentiates DMD iPSC-CMS from healthy control cells, consistent with the clinical Group 1 profile identified by Hathout, et al. (38). Only the biomarkers ANP32B, MK12 (MAPK12), troponin I (TNNI2, skeletal isoform), and fibrinogen/d-dimer did not demonstrate this pattern. The gene encoding ANP32B or acidic nuclear phosphoprotein 32, family member B is broadly expressed and implicated in signaling and growth (52). MK12 is involved in myogenesis and therefore may be a skeletal muscle specific response to stress (53). Troponin I (TNNI2) is produced almost exclusively in skeletal muscle (GTEX). Fibrinogen/d-dimer are produced exclusively by the liver (GTEX).

**Figure 7.**
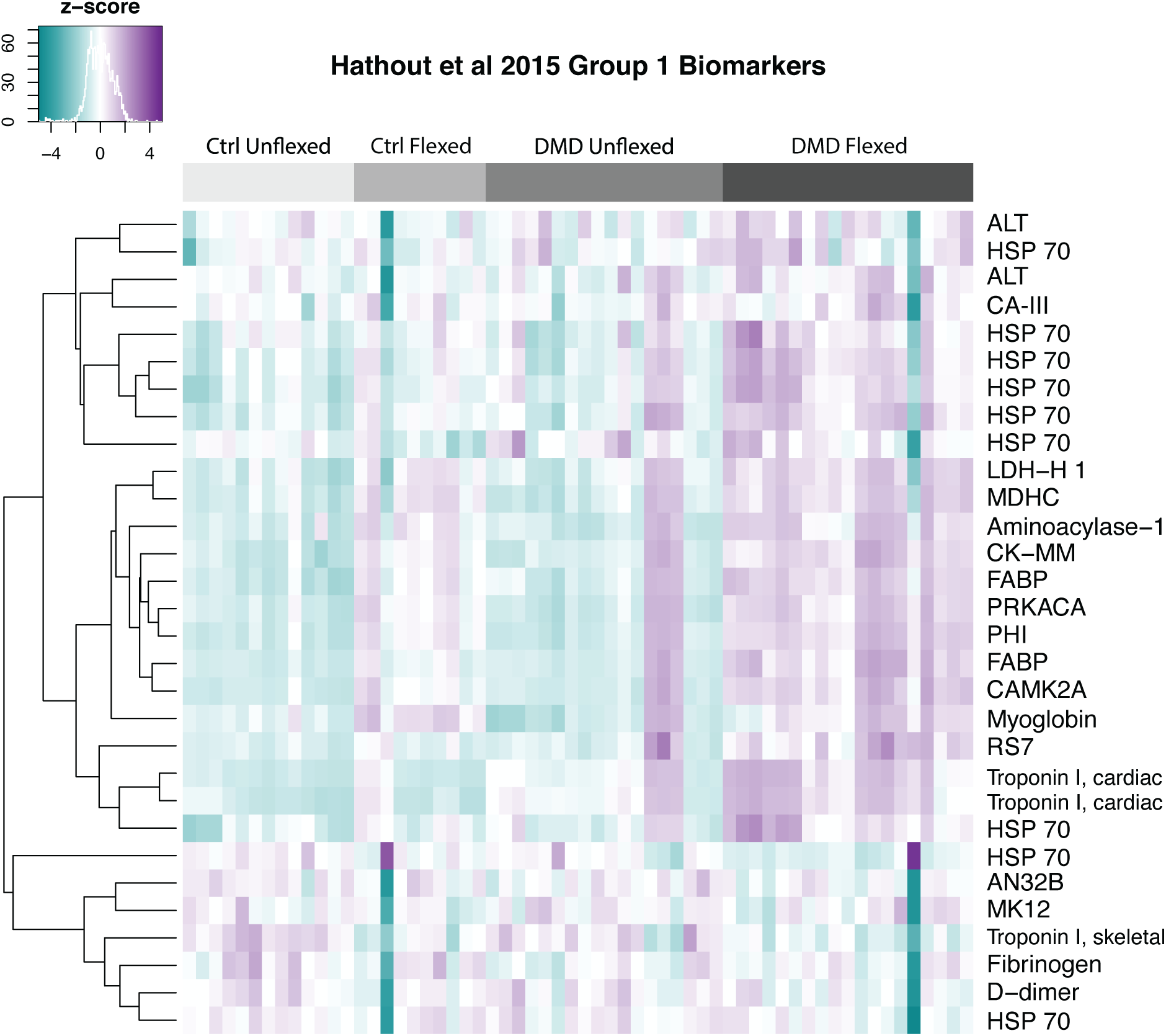
Biomarkers from DMD patients correlate with biomarkers released from flexed DMD iPSC-CMs. Hathout et al (38) conducted aptamer-based profiling on serum collected from DMD patients from multiple stages of disease progression. Group 1 markers are those seen in early DMD that are different from non-dystrophic controls. Group 1 markers decrease over the DMD lifespan, consistent with loss of muscle mass and replacement of muscle by fibrosis. In comparison to the aptamer-based protein biomarkers seen in DMD serum, media isolated from DMD iPSC-CMs subjected to equibiaxial strain showed similar elevated biomarkers, consistent with mechanical stress induced protein release from cultured cells. Several proteins are detected by multiple aptamers and are included for completeness.

## DISCUSSION

In vivo, cardiomyocytes are under constant cyclic stress due repetitive cardiac contraction. Membrane damage and repair are part of normal physiology, however certain diseases are associated with excessive membrane damage (54,55). Previous work has demonstrated in both the in vitro and in vivo setting that physiologic stress of the rat myocardium with isoproterenol induces transient membrane damage (55). Consistent with the role of dystrophin in skeletal muscle, the *mdx* mouse, which harbors a premature stop codon in *DMD* and is the most commonly studied DMD animal model, has demonstrated an increased susceptibility to cardiomyocyte membrane injury by an increase in afterload or treatment with isoproterenol (49). Collectively, the literature supports membrane fragility as the primary deficit in DMD and consequent membrane damage as the initial insult with a host of downstream consequences (11–13,21,22), and this is reflected by elevated serum proteins from both skeletal and cardiac origin in DMD patients (38,50).

Human iPSCs offer the advantage of harboring human mutations in its native cell context that can be differentiated and tested for treatment response (27,56). However, despite the ability to generate of iPSC-CMs, the conditions under which most cells are studied fail to simulate afterload and preload, and in the case of DMD cardiomyopathy, this is critical to creating micro-injury in the plasma membrane. Engineered heart tissues can be used to improve the maturity of iPSC-CMs enabling measurements of contractility; however, at present, methods for imparting dynamic mechanical stress are limited (28). In a recent report, Sewanan et and colleagues simulated pressure volume loops in decellularized porcine myocardium engineered heart tissue seeded with iPSC-CMs (57). By employing flexible membranes capable of deformation by equibiaxial strain, we successfully applied mechanical strain to iPSC-CMs in a physiologically meaningful way for the study of DMD-associated cardiomyopathy with clinically relevant protein biomarker outputs. We further validated protein release from stressed iPSC-CMs using an aptamer-based method to study thousands of proteins and directly comparing our results to protein profiling from human DMD serum. A striking correlation was seen when comparing a DMD patient data set to our cell-based model when 10% equibiaxial strain had been applied to the cells. iPSC-CMs cultured on these conditions are almost exclusively cardiomyocytes, and these cultures lack the typical infiltrative cells that characterize intact dystrophic heart or muscle. These data are consistent with notion that physiologic mechanical stress is necessary to bring out the clinically relevant phenotype in these cell models.

Several therapeutic approaches for the treatment of DMD have targeted increased membrane fragility. Poloxamer 188 is a triblock copolymer that has been extensively investigated for its membrane stabilization properties and has been shown to improve *mdx* hemodynamics and cardiomyocyte resistance to stretch-mediated injury (21,22). Poloxamer 188 is under clinical investigation (clinicaltrials.gov, NCT03558958) in a phase 2 study with the primary outcome of change in forced vital capacity, with secondary outcomes including cardiac endpoints. Enhancing native membrane repair is an alternative strategy. Mitsugumin53 (MG53) is a protein critical for muscle membrane repair that is also implicated in ischemic preconditioning (58,59). Recombinant MG53 has been shown enhance membrane repair and ameliorate aspects of muscle pathology in the *mdx* mouse (60). *Anxa6*, the gene encoding annexin A6 was discovered as a genetic modifier of muscular dystrophy, including genetic signals that implicated annexin A6 in cardiac function in a mouse model of muscular dystrophy (32). Overexpression of annexin A6 enhances membrane repair in skeletal myofibers, and exogeneously added recombinant annexin A6 similarly improved resealing of injured skeletal muscle myofibers (33,61). This work builds on those findings, demonstrating an ability to enhance membrane repair in injury that occurs in human DMD iPSC-CMs. Furthermore, recombinant annexin A6 also promoted resealing of healthy control iPSC-CMs, highlighting a conserved resealing mechanism in both normal and DMD cells.

Given that membrane damage is a part of normal physiology, endogenous repair mechanisms can be sufficient, provided injury is not so extensive. However, when faced with physiologic stress resulting in greater than normal membrane damage, as in the case of DMD, or stressors are greater than normal, as in a myocardial infarction, pathologic damage ensues. The schematic in **Figure 8** shows that recombinant annexin A6 enhances endogenous cardiomyocyte membrane repair processes. Given these findings, recombinant annexin A6 may be useful in treating genetic forms of cardiomyopathy that lead to increased baseline membrane fragility, as well as pathologic injury such as myocardial infarction or acute pressure overload where activation of membrane repair processes is essential for recovery from an acute insult. Additional studies will be required to confirm these findings in an in vivo format.

**Figure 8.**
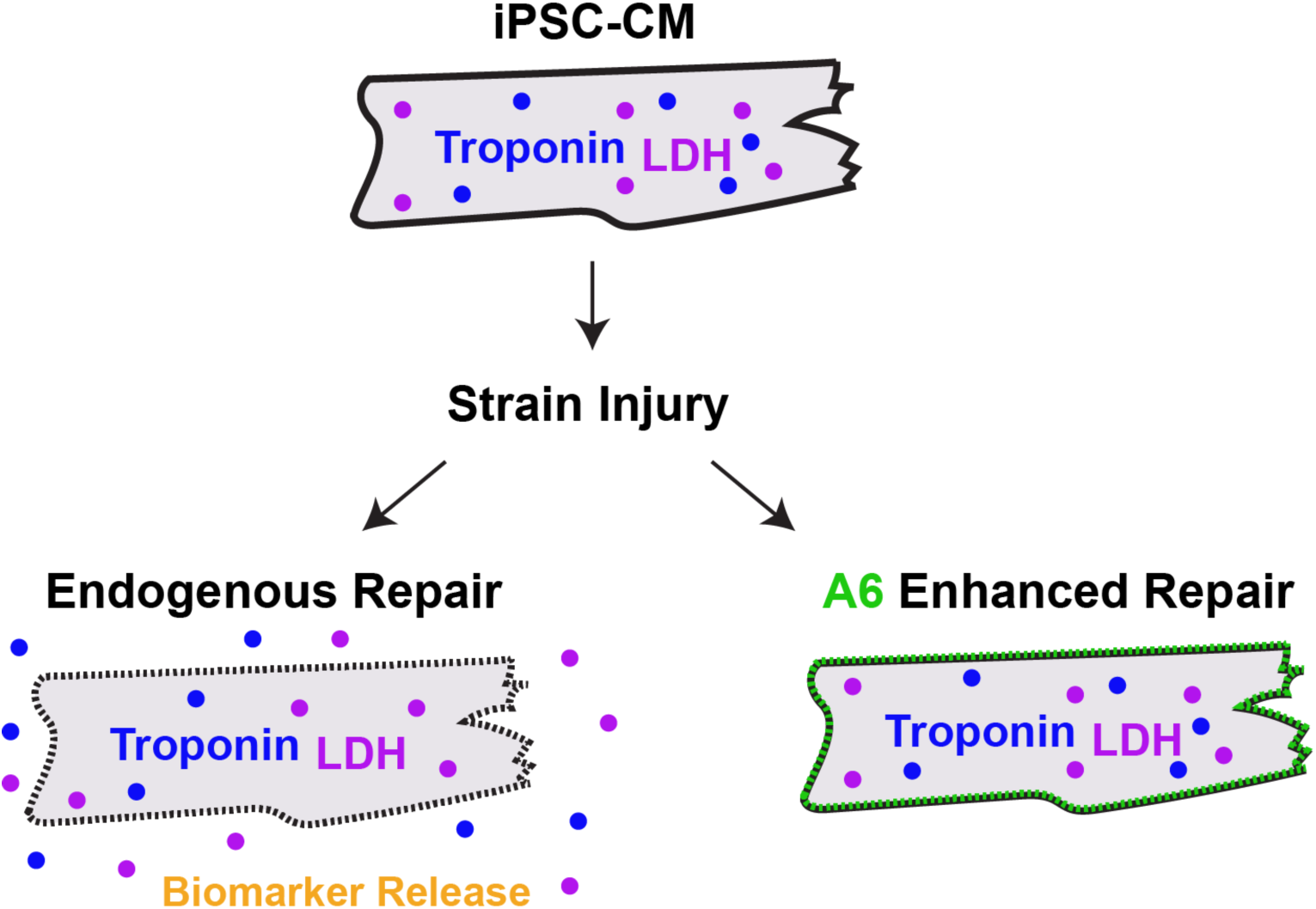
Recombinant annexin A6 promotes membrane repair in iPSC-CMs. Equibiaxial strain was employed to promote membrane injury in control and DMD iPSC-CMs. Recombinant annexin A6 promotes membrane repair as evidenced by decreased biomarker release in control and DMD iPSC-CMs.

### Conclusions and Limitations

In this work we applied equibiaxial strain to iPSC-CMs to assess the role of mechanical stress, and using this assay, we demonstrated a dose-dependent increase in protein biomarker release in DMD iPSC-CMs, as well as response to a protein resealing therapeutic. We also identified that proteins released into the media after equibiaxial strain reflected a similar profile to what is seen in early phase DMD patient serum, consistent with mechanical stress being an important driver of DMD pathology. While these data established the importance of incorporating mechanical stress into cell-based assays of cell injury, studies with additional cell lines may more appropriately reflect genotype-phenotype correlations for given *DMD* gene mutations. Similarly, it can be expected that healthy control lines also likely have a range of tolerance to injury, arising from both genetic and environmental conditions.

## Abbreviations

ACE: Angiotensin converting enzyme
DMD: Duchenne muscular dystrophy
iPSC: Induced pluripotent stem cell
iPSC-CMs: Induced pluripotent stem cell-derived cardiomyocytes
LDH: Lactate dehydrogenase
MG53: Mitsugumin53

## Acknowledgements

We thank the patient for donating cells for research.

